# Light modulates oscillatory alpha activity in the occipital cortex of totally visually blind individuals with intact non-visual photoreception

**DOI:** 10.1101/175851

**Authors:** Gilles Vandewalle, Markus J. van Ackeren, Véronique Daneault, Joseph T. Hull, Geneviève Albouy, Franco Lepore, Julien Doyon, Charles A. Czeisler, Marie Dumont, Julie Carrier, Steven W. Lockley, Olivier Collignon

## Abstract

The discovery of intrinsically photosensitive retinal ganglion cells (ipRGCs) marked a major shift in our understanding of how light is processed by the mammalian brain. These ipRGCs influence multiple functions not directly related to vision such as the synchronization of circadian rhythmicity, pupil constriction, the regulation of alertness and sleep, as well as the modulation of cognition. More recently, it was demonstrated that ipRGCs may also contribute to basic visual functions. The impact of ipRGCs on visual functions, independently of image forming photoreceptors, remains however difficult to isolate, particularly in humans. We previously showed that exposure to intense monochromatic blue light (465nm) induced awareness of light in a forced choice task in three rare totally visually blind individuals without detectable rod and cone function, but who retained non-visual responses to light, very likely via ipRGCs. The neural foundation of such light awareness in the absence of conscious vision is unknown, however. In this study, we characterized the brain activity of these three rare participants using electroencephalography (EEG), and demonstrate that unconsciously perceived light triggers an early and reliable transient desynchronization (i.e. decreased power) of the alpha EEG rhythm (8-14 Hz) over the occipital cortex. These results provide compelling insight into how ipRGC may contribute to transient changes in ongoing brain activity. They suggest that occipital alpha rhythm synchrony, which is typically linked to the visual system, is modulated by ipRGCs photoreception; a process that may contribute to the awareness of light in those blind individuals.

---

While vision is the best understood sensory system in the human brain, we are just beginning to understand the neuronal pathways mediating the ‘non-visual’ effects of ocular light exposure ^1,2^. Evidence for non-rod, non-cone retinal photoreceptors had been suspected in human ^3–6^ even before the discovery of intrinsically photosensitive retinal ganglion cells (ipRGCs) in animals ^7–9^. These retinal ganglion cells express the photosensory opsin melanopsin, which is maximally sensitive to short-wavelength blue light (∼480nm). IpRGCs are the primary photoreception channel mediating non-visual functions of light including entrainment of the circadian clock, suppression of the pineal hormone melatonin, pupillary constriction, and acute enhancement of alertness and cognitive brain functions ^1,10–12^. Melanopsin-driven intrinsic responses of ipRGCs are more sluggish than the response of classical photoreceptor. IpRGCs also receive incoming signals from rods and cones, however, such that their response output is time-locked to the photic stimulation ^1,13^. This photic response is then transferred to non-visual brain areas, including hypothalamic nuclei notably involved in circadian rhythmicity, and sleep-wake regulation ^14,15^. IpRGC signals also reaches parts of brain primarily mediating vision, such as the lateral geniculate nucleus of the thalamus ^13,15^ and studies suggest that ipRGCs contribute to crude vision, brightness detection, and adjustment of rod and cones sensitivities ^16–21^. Similarly to animal models, many human non-visual responses to light are more sensitive to shorter wavelength (blue) light exposures making them likely to be under ipRGC influence ^5,22–26^. Isolating ipRGCs and melanopsin roles from those of rods, cones and other RGCs can be achieved in animals by knocking down, amplifying, or modifying part of the photoreception system ^19,27,28^. The exact role of ipRGCs remains difficult to isolate in humans, however, where specific genotypes perturbing the different parts of the retinal photoreception system are not available. Recently, new psychophysical technique has been developed in order to target melanopsin separately from the cones ^17,25,29–31^.

To study ipRGCs in isolation, we capitalize on a rare and small group of totally visually blind human individuals-a dozen-have been reported to exhibit normal 24-hour circadian rhythms and have intact melatonin suppression, pupil constriction and/or circadian phase resetting in response to nighttime ocular light exposure ^3,32– 34^. Critically, these light effects did not occur when these participants had their eyes covered. Neuro-ophthalmologic exam confirmed absence of rod and cone function in these individuals, strongly supporting that these rare individuals likely retain light perception via ipRGCs. This hypothesis is further supported by the observation that shorter wavelength light, in the blue range (∼ = 470 nm) was shown to be more efficient in triggering an alerting or pupil response in such individuals ^33,34^. Furthermore, when exposed to blue light during a cognitively challenging task as compared to while in darkness, these atypical blind individuals showed enhanced activation in a widespread subcortical and cortical network, including the occipital cortex ^35^. Altogether, these results support the notion that these rare blind individuals provide a unique human model system to study the role of ipRGCs in the absence of conscious vision.

At the frontier between vision and non-vision, these rare patients present the ability to non-randomly guess the presence or absence of light in a two-alternative choice forced task (2AFC) indicating some sort of awareness for light in the absence of conscious vision [n = 1 ^33^; n = 3 ^35^]. The brain mechanisms of such awareness of light are unknown. Cortical oscillations in the alpha range (8-14 Hz) may provide important insights to address this issue. One the one hand, exposure to blue light at night *increases* high alpha power in sighted individuals (∼10-12Hz) ^24,36,37^, and in a blind individual with presumably intact ipRGCs (∼8-10 Hz) ^33^. These *sustained* changes in alpha power are typically reported over the *entire* cortex and following prolonged exposure to light; they seem to take minutes to tens of minutes to be detected. They have not been investigated, however, at shorter time scales using exposure to lights lasting only of seconds. On the other hand, alpha oscillations in sighted individuals show transient responses to visual stimulation, which emerges in the order of milliseconds to seconds. These are thought to reflect changes in visual attention allocation and are typically observed *locally* as a *transient decrease* in alpha power, or a transient desynchronization, over the area of the brain that processes the stimulus presented ^38–42^. Interestingly, recent evidence suggests that these more transient oscillatory changes could be modulated by prior exposure to bright blue light, likely implicating ipRGCs ^43,44^. However, the transient response to light has not been investigated in blind participants retaining non-visual responses. Since awareness for light appears somewhat typically visual, we hypothesize that exposure to light would transiently decrease alpha power over the occipital cortex during light exposure in these rare blind individuals, supporting that light awareness arises from a specific impact on visual attention.

To test this hypothesis, we analyzed the EEG recordings acquired while three totally visually blind individuals with intact non-visual responses to light participated during a forced choice task for which they choose the presence or absence of light non-randomly despite their complete lack of conscious vision ^35^. Spectral analysis and EEG source reconstruction computations were performed to quantify short-term (10s) light-induced changes in alpha rhythm over the occipital cortex task.

## Results

EEG was recorded during forty 20s-trials of an AFCT during which blue light [9.7 × 10^14^ photons/cm^2^/s; peak = 465nm; Full Width at Half Maximum (FWHM) = 27nm] was pseudo-randomly presented during the first or last 10s (10s light ON and OFF). As reported previously ^35^, both participant 1 and 2 showed a very high accuracy in guessing when the blue light was presented (95%, and 80%, both *p*<.001). Participant 3 also showed performance rates different from chance although accuracy was inverted (30%, *p*=.008). EEG spectral analysis of the alpha power difference between the light-ON *versus* light-OFF across occipital electrodes and over the entire 10s time windows revealed significantly reduced alpha power in all three participants (participant 1: *p*=.034; participant 2: *p*=.024; participant 3: *p*=.028) (Fig. 1A). This sustained difference between conditions was strongest in the upper alpha range [11-14Hz] for each of the three participants (participant 1: 12.1-12.8Hz; participant 2: 12.8-13.9Hz; participant 3: 11.3-12.5Hz)

**Figure 1.**
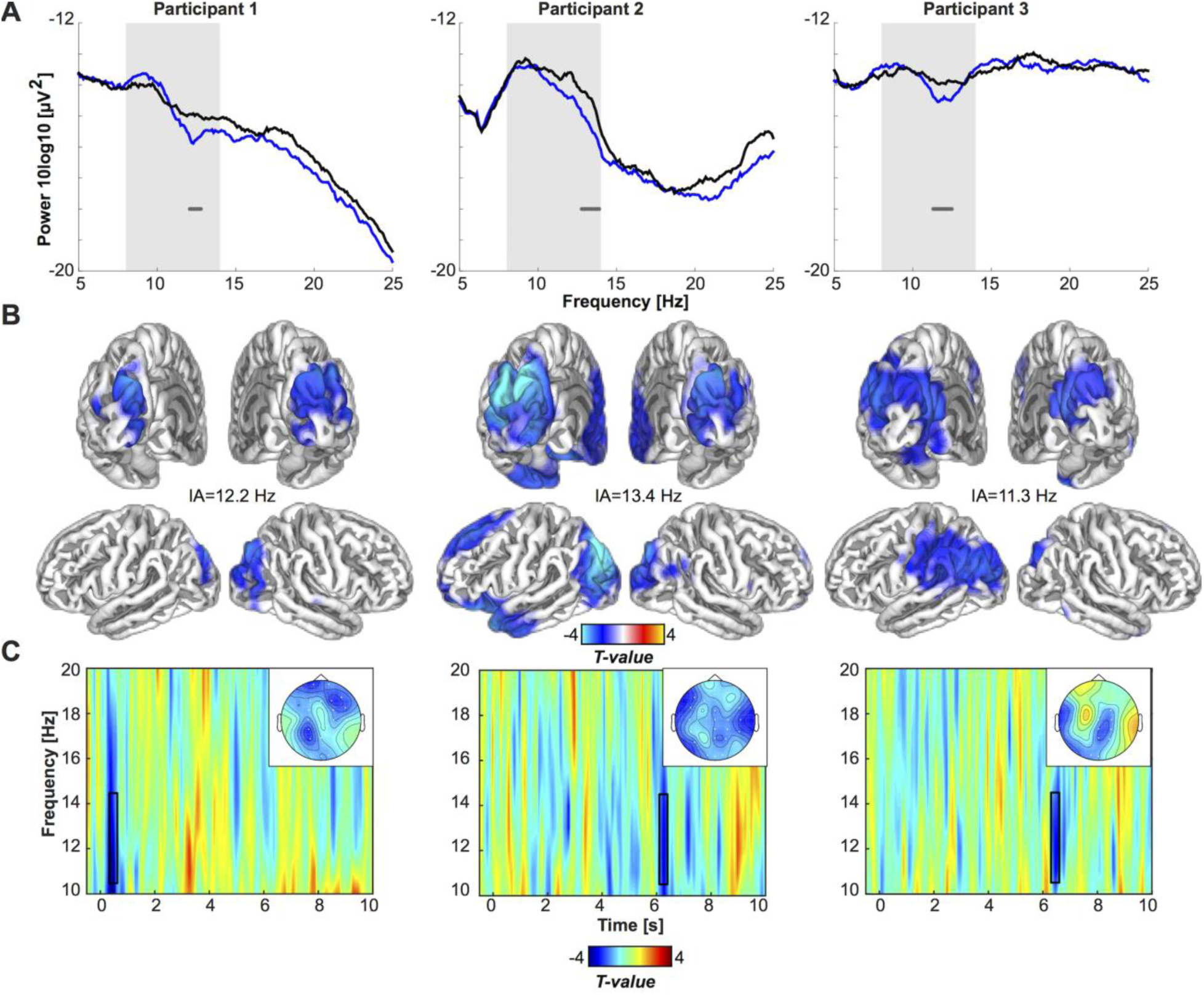
Desynchronization of the occipital alpha/low beta rhythm during a 10 second of stimulation with blue light in three blind participants with intact non-visual photoreception. **(A).** Frequency spectra for the entire 10s period of blue light (blue), and darkness (black) show differences between conditions in all three participants (horizontal gray bar, p<.05, permutation-cluster-corrected for multiple comparisons) for the frequency band of interest (shaded area, 8-14Hz). **(B).** Source reconstructions depict the difference between conditions (thresholded at p<.05, FDR corrected for multiple comparisons), at the individual alpha frequency showing greatest light-induced decrease (see main text). Statistical maps (t-values) of the topographies with the absolute difference between conditions at the individual alpha frequencies show that the spatial extent of the decrease in alpha power is largely restricted to occipital electrodes. **(C)** Time-frequency representations illustrate differences between the light-ON versus light-OFF condition in time. Open rectangles highlight the time range of greatest difference (*p*<.05, FDR-cluster-corrected for multiple comparisons)

Subsequent source reconstruction was performed (i.e. not restricting to occipital electrodes) on the entire 10s time window, using the individual peak in alpha power difference between light-ON and light-OFF (participant 1: 12.2 Hz; participant 2: 13.4 Hz; and participant 3: 11.3Hz; see highlighted peaks on Fig. 1A). Although the analyses were not restricted to the posterior part of the brain, the neural generators of the alpha power decrease during light-ON was localized primarily to occipital cortex in participants 1 and 2, and to occipito-parietal cortex in participant 3 (Fig. 1B). In addition, a distinct conjunction analysis, seeking for significant sources common to all participants combining all three statistical maps, revealed an area in bilateral superior occipital gyrus (SOG) (Fig 2). The light-induced decrease in alpha power observed over occipital sensors seems therefore to be mostly confined to the occipital cortical areas.

**Figure 2.**
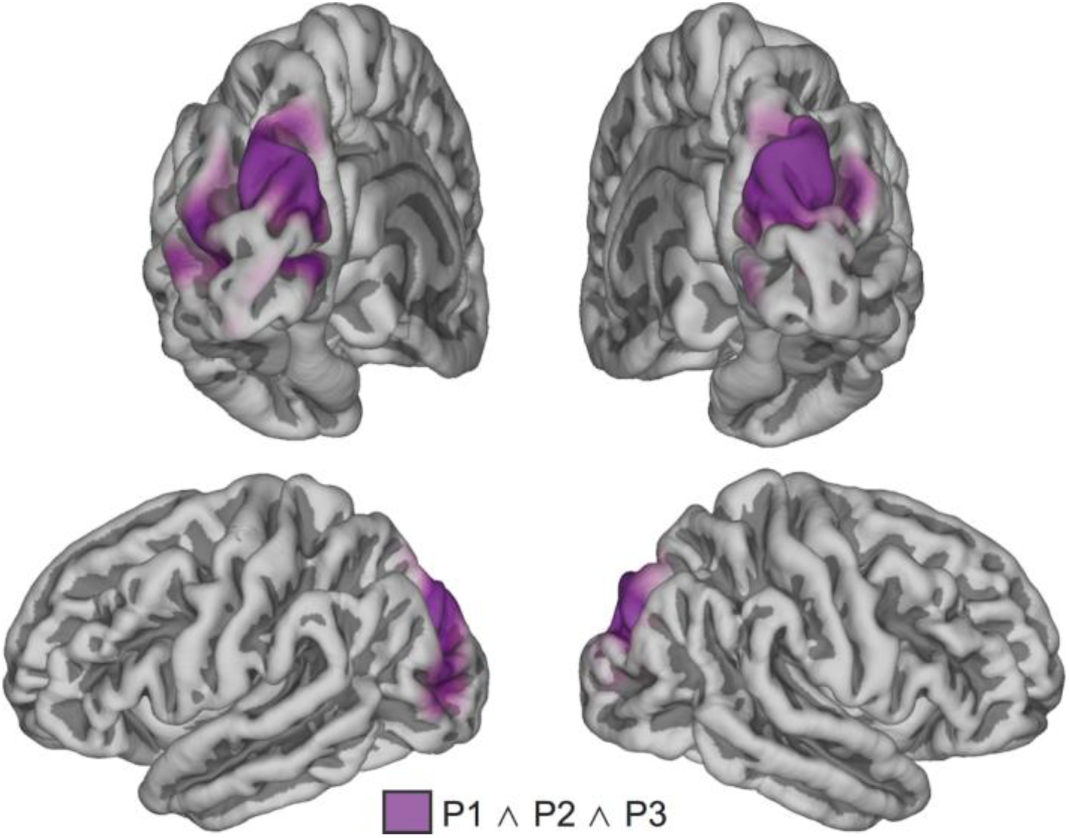
Conjunction analysis across the source statistical maps identified an area in bilateral superior occipital gyrus and cuneus as a significant (p<.05) cortical source of the alpha suppression in response to blue light common to all three participants.

Finally, Figure 1C shows the time-frequency maps for the statistical difference between the light-ON and light-OFF condition for each of the three participants. This exploratory follow-up and distinct analyses on the upper alpha range (11-14Hz) performed across time and space revealed transient alpha suppression effect. These transient statistically significant changes indicate the time points at which the sustained light-induced reduction in alpha power detected across the 10s exposure is strongest. While participant 1 showed a broadband significant desynchronization shortly following stimulation onset (400-600ms, *p*=.01), participant 2 (6200-6400ms, *p*=.012) and 3 (6400-6600ms, *p*=.038) showed significant desynchronization only after several seconds.

## Discussion

Some rare totally blind individuals with preserved non-image-forming photoreception can guess successfully on the presence or absence of light in a forced choice task ^33,35^. In this study, we investigated the brain mechanisms underlying this awareness for light using EEG data recording during the 2AFC guessing task. We show that 10s of light exposure is able to reliably induce a transient and local decrease of alpha power over the occipital cortex in the three participants tested. Although the exact frequency seems to slightly vary across individuals (peak of alpha power decrease between 11.3Hz and 13.4 Hz), this decrease is more prominent in the higher alpha band between 11 and 14Hz rather than in the lower alpha.

Transient alpha oscillations are thought to reflect a sensory gating, or prioritization, mechanism ^40,41^. For example, fluctuations in the cortical alpha rhythm have been associated with visual-spatial, sensory, and object-based attention ^39,45–47^, including response to blue-enriched white light ^43^. Specifically, transient reduced alpha power typically reflects enhanced neuronal activity, or excitability ^48–50^, while transient enhanced alpha power is considered a marker for cortical inhibition ^40,41^. For example, if alpha power in a given sensory area is low, participants are more likely to perceive a near-threshold event in the same modality ^49,51,52^. If transient reduced alpha power is a marker for increased cortical excitability in visual areas, or indeed a precursor to visual awareness, this could explain why participants made non-random selections about the presence or absence of the blue light during the 2AFC task.

In our view, the source of our effects is most parsimoniously explained by ipRGCs photoreception because (1) these cells are known to be preserved in individuals with outer retinal degeneration ^53^, (2) ophthalmological examination confirmed atrophy of the retinal pigment epithelium and found no detectable functional responses from rods and cones ^33^, (3) light effects were more pronounced using 460-to 480-nm monochromatic light as compared with other wavelengths ^33^, and (4) recent data show that the dynamics of pupillary constriction in a blind human with outer retinal degeneration was compatible with the exclusive involvement of ipRGCs ^34^. Moreover, participants had no conscious light perception and no visually evoked EEG responses to 800 flashes of light. Including other wavelengths of light and showing that EEG modulation was stronger with exposure around 480 nm could have strengthen our finding, but it would not add to the fact that light modulated EEG activity in individual with no detectable residual outer retina function. Therefore, although we cannot formally exclude a marginal contribution of remaining but yet undetected rod or cones, we provide compelling evidence that the melanopsin-driven photosensitivity of ipRGCs feed to the human visual cortex and trigger some awareness for light.

This assumption is in fact supported by recent data in sighted individuals exposed to metameric light that stimulated melanospin while leaving the other retinal photoreceptor equally stimulated (such that the appearing color of the light remains almost constant). This type of light stimulation has been used to show that melanopsin-driven ipRGCs output modulate activity of the frontal eye field, a brain region involved in visual attention and ocular motor responses ^31^ and of the occipital cortex ^29^, extending previous animal research ^54^. In addition metameric light geared toward melanopsin indicated that ipRGCs output contribute to conscious visual perception of brightness in human ^17,29^, but only if the modulation of light spectral composition was relatively slow ^29,30^. One could assume that, in blind individuals with intact ipRGCs, brightness is a difficult concept to elaborate on that can be detected in the conscious percept through a forced guess. If true, then this could mean that brightness detection in sighted individual is, at least in part, mediated through a transient change in alpha oscillation over the occipital cortex. This remains to be investigated. Alternatively, the moment of most alpha desynchronization could correspond to the average moment at which each participants decides on his/her guess based on brightness perception.

The significant transient change in alpha occipital activity appears to be brief in the time-frequency analysis (∼200ms). This may be emphasized by the stringent statistic we apply to our small sample size. Indeed, simple spectral analysis yielded significant alpha reduction on average over the entire 10s trials. One should consider the time-frequency as an indicator of the moment at which desynchronization is strongest. This moment varied from ∼500ms in participant 1 to ∼6s in the other two participants, which is long, in any case, with respect to classical rod-cone photoreception. This relatively late onset response may be related to the relatively sluggish responses of the intrinsic melanopsin-driven light sensitivity of ipRGC in the absence of classical photoreceptor inputs ^1,8,13^.

IpRGCs are considered to mediate at least in part the impact of light on alertness and attention ^22^, which is evident through a sustained and global decrease in delta (0.5-4Hz) and theta (4-8 Hz) power in EEG recordings of spontaneous (i.e. task free) brain activity ^36,55^ concomitant to an increase in alpha power ^24^. In situation of vigilance (like in the Zaidi’s paper, 2007), the impact of light would express at late temporal scale with an increase in alpha power. In contrast, when attending transient stimuli changes, light appears to induce a transient reduction of alpha power at relatively short time-scale. These results therefore bring new information about how ipRGC can play a prominent role on visual functions that is distinct from the slow fluctuating effects observed in relation to alertness circuits. How related are the perceptual role of ipRGCs and their impact on vigilance and attention requires further investigation.

Finally, the fact that oscillatory changes in the current study seem to be largely confined to occipital areas, as well as the observation that participants demonstrate non-conscious behavioral sensitivity to the presence of blue light could have profound consequences for cortical reorganization of occipital cortex in blind participants with intact ipRGCs. Specifically, the presence of residual pathways still attuned to ipRGCs influence in the occipital cortex of blind people could prevent the full reorganization of those regions toward non-visual functions typically observed in totally blind people ^56^, and therefore might weight in the decision to attempt sight restoration in this population^57,58^.

In summary, we demonstrate that short exposure to light, in the order of several seconds, reduces alpha power in occipital cortex in three blind human participants which present an awareness for the presence of light. This effect is likely attributed to ipRGCs and could be related to their suggested role in brightness detection in sighted individuals ^17,29^. Using a human model of isolated ipRGCs photoreception has proven to be insightful to better apprehend melatonin suppression, pupil and cognition regulation, and circadian phase resetting by light ^3,32– 35^. The present results extend these previous results by strengthening the view that ipRGCs photoreception influence the visual system to a much greater extent than previously thought.

## Methods

Except for electrophysiological data analyses, methods included here were described previously ^35^.

### Participants

Participants were three totally visually blind individuals [1 female, range 60-67 years; see table 1 for detailed characteristics, previously reported in ^35^]. Although none of the participants reported any conscious awareness of light, an intact melatonin suppression responses to white or blue light was previously demonstrated in all three ^32,33^. All three participants had had previous standard ophthalmologic examination in Boston include standard VEP or electroretinogram (ERG). Participant 1 had pupil muscle damage during a surgery, while participant 3 had no clearly distinguishable pupil. Only participant 2 exhibited pupil constriction if a pen-light exposure was continued for more than a few seconds ^34^. In previous visits to Boston, a fundoscopic examination in Participants 1 and 2 confirmed atrophy of the retinal pigment epithelium, with thinning of retinal vessels and bone spicule pigmentation. These findings were confirmed on several occasions by different ophthalmologists who examined the participants independently.

### Experimental design

The experiment was conducted in the afternoon in an electromagnetically shielded and sound attenuated room in complete darkness. Visual stimulation was elicited via an array of 48 blue light emitting diodes [LED, peak = 465nm, Full Width at Half Maximum (FWHM) = 27nm] behind a 21×11 cm diffusion glass with ultraviolet and infrared filters. Photon flux at corneal level was 9.7 × 10^14^ photons/cm^2^/sec. Brief blue-light flashes were administered during four blocks of 200 trials (duration=500ms, ISI=1600-1800ms), totaling 800 trials, to record potential visually evoked potential (VEP). No EEG responses could be detected further reinforcing that no residual rod and cone function remained [see ^35^]. EEG recordings were also acquired during a two-alternative forced choice task (2AFC). Each trial lasted 20s during which blue light was pseudo-randomly presented during the first or last 10s (10s light ON and OFF). The halfway point of the trial was signaled by a brief auditory pure tone (500ms, 1000Hz). After each trial, the participant was asked whether the photic stimulation occurred during the first or second half of the trial. Participants were presented with 40 trials in total resulting in 80 segments, half with and half without light stimulation. Due to a scripting error, only the overall performance on the task is available for participant 1, i.e., responses to individual trails were not recorded. Actually, this participant provided the most numerous correct “guesses” about light-presence (95% of correct guesses) meaning that he would have had only 2 incorrected guesses to contrast with the 38 correct ones, precluding any reliable analyses between EEG and behavioral measures at the individual trial level.

### EEG data acquisition and preprocessing

EEG was recorded continuously from 40 Ag-AgCl electrodes on an extended 10-20 system (Neurosoft Inc., Sterling, VA), using a sampling rate of 1000Hz. The signal was filtered online using a band-pass filter (0.1-100Hz), and referenced to the algebraic average of the two mastoids. Impedances were kept below 5 kΩ. The signal was filtered offline with a low-pass filter with a cut-off at 30Hz, re-referenced to the algebraic average across electrodes, and downsampled to 150Hz. Due to the long duration and limited number of trials of the 2AFC, extended infomax independent component analysis (ICA) was performed with a weight change stop criterion of 10^−7^. Components reflecting ocular artifacts (e.g. eye blinks and lateral saccades) were rejected based on visual inspection of topography and time course.

### Spectral analysis

All offline analysis was conducted using Fieldtrip ^59^, under Matlab (The Mathworks Inc, Natick, MA). Analysis was performed separately for the 10s exposure to light (Light ON), and darkness (light OFF). Spectral analysis was applied over the entire 10s period for frequencies between 5 and 25Hz, using the multitaper method ^60^ with 19 Slepian tapers, resulting in a frequency smoothing of ±1Hz. Statistical analysis was performed on the log-transformed power spectra, and centered on the alpha band (8-14Hz) in occipital electrodes (O1, Oz, and O2). Between-trial comparisons for the light-ON *versus* light-OFF segments were conducted for each individual participant using independent sample t-tests. A cluster permutation approach was used (500 permutations) to control for multiple comparisons using a maximum cluster extent threshold ^61^. Permutation tests provide a robust inferential statistic, without assumptions about the shape of the sampling distribution. Furthermore, by selecting a liberal alpha-range (8-14Hz), the method optimally captures potential variation in the individual alpha peak frequency. To explore the time-course of alpha power modulation in response to light further, time-frequency analysis was performed using 6 cycle Morlet wavelets (100ms steps, 1Hz frequency resolution). Here, the frequency-averaged alpha power time course was compared between the light-ON versus light-OFF condition and clustering was performed across time (0-10s post stimulus onset) and space (electrodes). We want to emphasize that this test followed the initial comparison of the frequency spectra, and should therefore be considered exploratory.

### Source-level analysis

In line with the recommendations by Gross and collaborators ^62^, statistical analysis of the differences between the light-ON and light-OFF condition were performed at the sensor level, and subsequent source reconstruction was performed to identify the generators. Source reconstruction was performed using structural MR-images from each of the three blind participants. T1 weighted images (voxel size: 1×1×1mm, TR=2.3s, TE=2.91ms, flip angle=9°, field of view 265×224mm) were acquired on a 3-T TIM-TRIO MR scanner (Siemens, Erlangen, Germany). Individual T1 MRI images were segmented in cerebrospinal, white matter, grey matter and skull compartments. The white and grey matter brain compartments were used to construct individual head models using the Boundary Element Method (BEM) ^63^. Source activity was projected on a regular three dimensional grid (10×10×10mm), using a frequency domain beamformer, called Dynamic Imaging of Coherent Sources ^64,65^. A regularization of 15% was applied. To identify robust peak differences in the alpha range, statistical maps for the difference between the light-ON versus light-OFF conditions were computed using independent-sample t-tests, using a permutation approach, with a threshold of α = 0.05 (FDR corrected).

## Acknowledgements

We thank Amélie Apinis-Deshaies, Hélène Blais, Nathalie Bouloute, Annick Cartier, André Cyr, Stéphane Denis, Mathieu Desrosiers, Sonia Frenette, Simon Girard, Philippe Grenier-Vallée, Carollyn Hurst, Marjolaine Lafortune, Frédéric Lesage, Nicolas Ouakli, Caroline Reinhardt, Manon Robert, Zoran Sekerovic for their invaluable help. We also thank Dr. Joseph F. Rizzo III, Dr. Eliot L Berson, and Dr. Keith H. Chiappa for performing initial neuroophthalmology examinations of the patients and Dr. Elizabeth B. Klerman for medical supervision of the studies conducted in Boston. This study was partly supported by the Réseau Vision du Québec and Réseau de Bioimagerie du Québec (RBiQ), the Fonds de Recherche en Santé du Québec (FRSQ) and a European Research Council starting grant attributed to OC (MADVIS: ERC-StG 337573). GV and OC are supported by the Fonds National de la Recherche Scientifique (FNRS, Belgium) Initial screening and melatonin suppression confirmation was supported by the National Institute of Neurological Disorders and Stroke (R01NS40982 to CAC and SWL), The Wellcome Trust, UK (060018/B/99/Z to SWL), and the National Center for Research Resources through the Harvard Clinical and Translational Science Center at Brigham and Women’s Hospital (M01RR02635 and UL1RR025758). JTH was supported in-part by a Cephalon Clinical Fellowship in Circadian Medicine and a National Heart, Lung and Blood Institute fellowship in Sleep, Circadian and Respiratory Neurobiology (T32 HL079010), Division of Sleep Medicine, Harvard Medical School.

## Author contributions

GV, OC, JC, SWL, JTH, designed the experiment; GV, OC, VD, GA, performed the experiment; CAC, JTH, SWL identified and recruited the rare participants included in the study; MJVA analyzed the data in interaction with GV and OC; FL, JD, MD, contributed to study design and statistical analyses; MJVA, GV, OC, JC, SWL, VD wrote the manuscript, JTH, GA, FL, JD, CAC, MD edited the manuscript.

---

**SUPPLEMENTARY TABLE S1:**
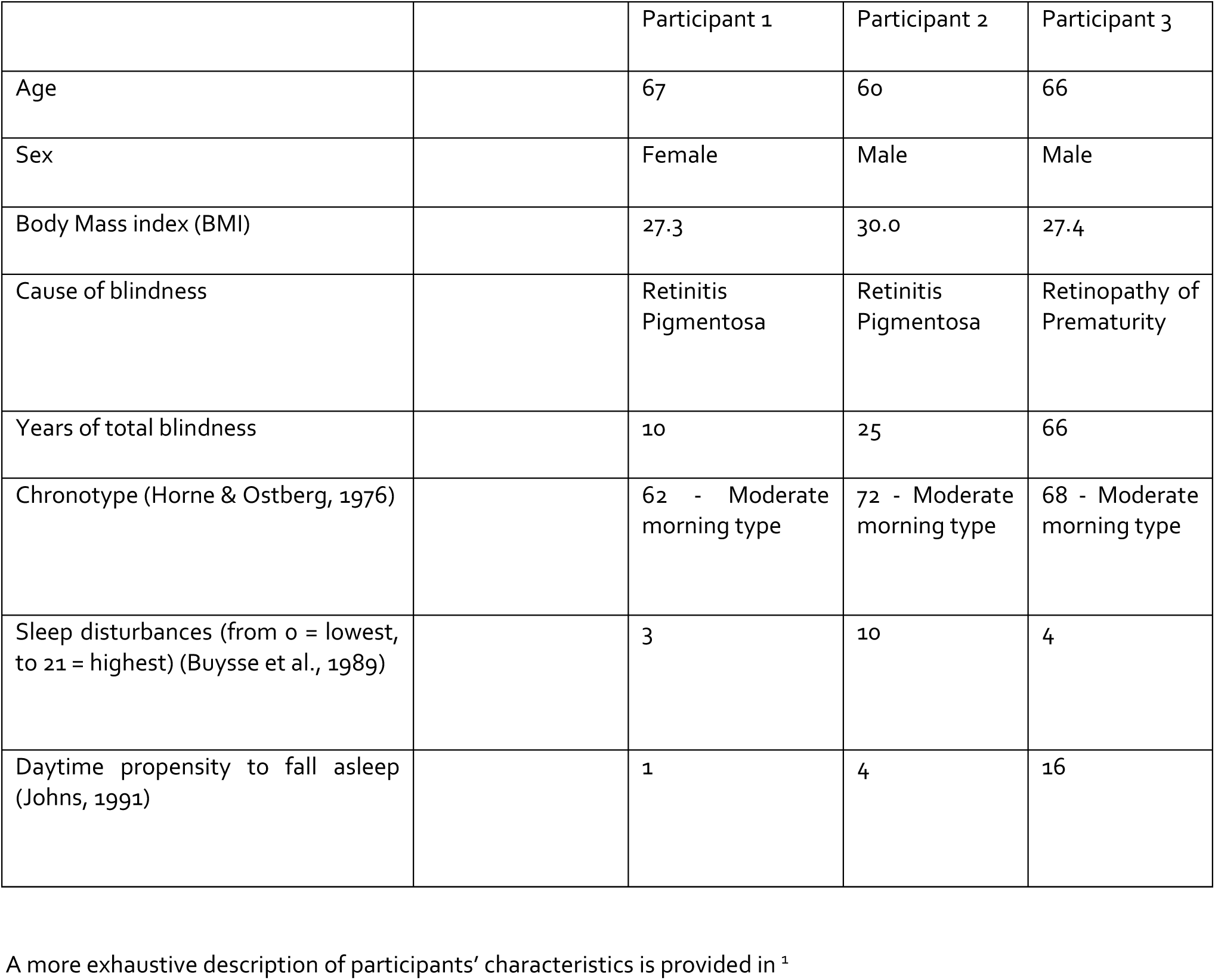
Participants characteristics

|  |  | Participant 1 | Participant 2 | Participant 3 |
| --- | --- | --- | --- | --- |
| Age |  | 67 | 60 | 66 |
| Sex |  | Female | Male | Male |
| Body Mass index (BMI) |  | 27.3 | 30.0 | 27.4 |
| Cause of blindness |  | Retinitis Pigmentosa | Retinitis Pigmentosa | Retinopathy of Prematurity |
| Years of total blindness |  | 10 | 25 | 66 |
| Chronotype (Horne & Ostberg, 1976) |  | 62 - Moderate morning type | 72 - Moderate morning type | 68 - Moderate morning type |
| Sleep disturbances (from 0 = lowest, to 21 = highest) (Buysse et al., 1989) |  | 3 | 10 | 4 |
| Daytime propensity to fall asleep (Johns, 1991) |  | 1 | 4 | 16 |

A more exhaustive description of participants' characteristics is provided in <sup>1</sup>

**Supplementary Figures S1.**
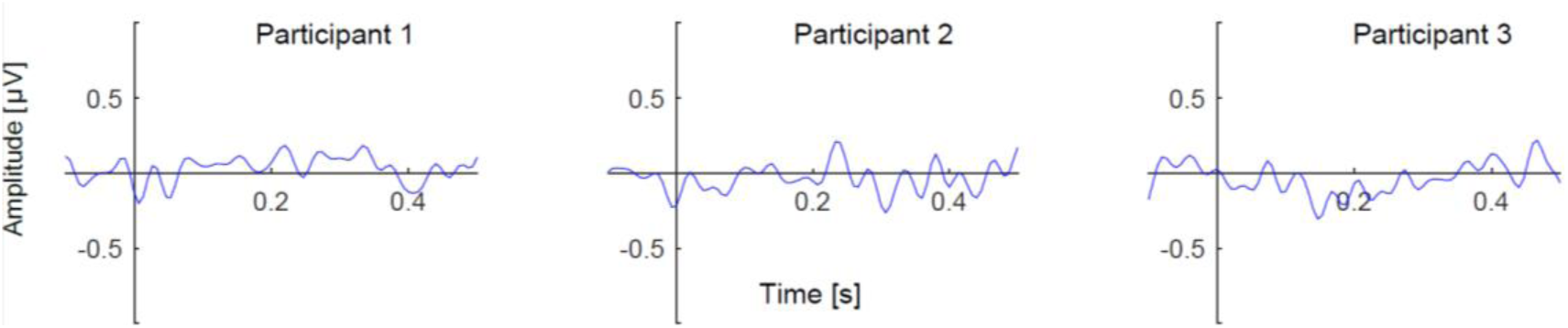
Trial averages in the time domain for brief photic stimulation with blue light, relative to baseline showing a marked absence of visual evoked potentials (VEP; 800 trials).

